# Reward signalling in brainstem nuclei under glycemic flux

**DOI:** 10.1101/243006

**Authors:** Tobias Morville, Kristoffer Madsen, Hartwig R. Siebner, Oliver J. Hulme

## Abstract

Phasic dopamine release from mid-brain dopaminergic neurons signals errors of reward prediction (RPE). If reward maximisation is to maintain homeostasis, then the value of primary rewards should be coupled to the homeostatic errors they remediate. This leads to the prediction that RPE signals should be configured as a function of homeostatic state and thus, diminish with the attenuation of homeostatic error. To test this hypothesis, we collected a large volume of functional MRI data from five human volunteers on four separate days. After fasting for 12 hours, subjects consumed preloads that differed in glucose concentration. Participants then underwent a Pavlovian cue-conditioning paradigm in which the colour of a fixation-cross was stochastically associated with the delivery of water or glucose via a gustometer. This design afforded computation of RPE separately for better- and worse-than expected outcomes during ascending and descending trajectories of physiological serum glucose fluctuations. In the parabrachial nuclei, variations in regional activity coding positive RPEs scaled positively with serum glucose for ascending and descending glucose levels. The ventral tegmental area and substantia nigra became more sensitive to negative RPEs when glucose levels were ascending. Together, the results show that RPE signals in key brainstem structures are modulated by homeostatic trajectories of naturally occurring glycemic flux, revealing a tight interplay between homeostatic state and the neural encoding of primary reward in the human brain.

## Introduction

A basic assumption of many models of adaptive behavior, is that the value of primary rewards are modulated by their capacity to rectify future homeostatic deficits (Pompilio et al. 2006; Cabanac 1971). Compatible with this notion, deprivation-induced hypoglycaemia increases willingness to work for food in rats (Sclafani et al. 1970), in humans (Pelchat 2009), as well as the subjectively reported pleasure (Cabanac 1971). Catecholamine dopamine is a neurotransmitter that plays a key role in signalling reward (Haber & Knutson 2010) and is involved in behavioural reinforcement, learning and motivation (Berridge 2006; Schultz et al. 1997). Via meso-cortical and mesolimbic dopaminergic projections, synaptic dopamine release modulates the plasticity of cortico-striatal networks and hereby sculpts behavioural policies according to their reward contingencies (Haber & Knutson 2010; Schultz 2015). Patterns of phasic dopaminergic firing have been demonstrated to follow closely the principles of reinforcement learning, encoding the errors in the prediction of reward (O’Doherty et al. 2004; Schultz et al. 1997; Rangel et al. 2008; Tobler et al. 2005). Reward prediction error (RPE) signals are commensurate with the economic construct of marginal utility, defined as the additional utility obtained through additional units of consumption, where utility is a subjective value inferred from choice (Stauffer et al. 2014; Schultz 2005; Schultz 2015).

Although animals are motivated by a homeostatic deficit of thirst or hunger, homeostatic states are rarely considered as relevant modulators of dopaminergic signalling of reward prediction errors. In typical paradigms involving cumulative consumption, the homeostatic deficit gradually diminishes as the animal plays for consumption of water or sugar-containing juice. Eventually, the animal rejects further play, presumably because the marginal utility of consumption diminished to a point of indifference. Interestingly, a recent electrophysiology study in rats, demonstrated that oral consumption of sodium solution causes phasic dopaminergic signals in the nucleus accumbens, that are modulated by sodium depletion (J. J. Cone et al. 2016).

There is now growing evidence for a multifaceted interface between dopamine mediated reward-signalling and the systems underpinning energy homeostasis. Firstly, dopamine neurons in the ventral tegmental area (VTA) express a suite of receptors targeted by energy-reporting hormones ghrelin, insulin, amylin, leptin and Glucagon Like Peptide 1 (GLP-1, Ferrario et al. 2016; Palmiter 2007). This provides numerous degrees of freedom for flexibly interfacing between homeostatic state and reward signalling. Although hormonal modulations of phasic dopamine are yet to be fully scrutinised, there is emerging evidence that circulating factors do indeed modulate its magnitude. For instance, amylin, a hormone co-released with insulin, acts on the VTA to reduce phasic dopamine release in its mesolimbic projection sites (Mietlicki-Baase et al. 2015). In terms of neuronal input, there are many such opportunities for the appetitive control of dopamine mediated signalling.

Appetitive control can be delineated into three interacting valuation systems (Sternson & Eiselt 2017). The first system generates a negative valence signal which involves activity of the Agouti-related peptide (AgRP) neurons of the arcuate nucleus of the hypothalamus (ARC). Activity of ARC_AgRP_ neurons reports on energy deficits, inhibits energy expenditure, and regulates glucose metabolism (e.g. Aponte et al. 2011; Dietrich et al. 2015; Luquet et al. 2005; Cansell et al. 2012). ARC neurons that contain peptide products of pro-opiomelanocortin (POMC) form an opponent code compared with ARC_AgRP_ neurons. The balance between the two neuronal ARC sub-populations putatively encodes the value of near-term energetic states, becoming rapidly modulated just prior to food consumption (Mandelblat-Cerf et al. 2015). The second system codes positive valence signals and consists of circuits involving the lateral hypothalamus (LH). It is linked to positively reinforcing consummatory behaviours via its dopaminergic projections, assumed to trigger positive feedback to keep consumption going during feeding bouts. The third valuation system involves calcitonin gene-related protein (CGRP)-expressing neurons in the (PBN) that potently suppress eating when activated, but do not increase food intake when inhibited. PBNcgrp neurons are activated by signals associated with food intake, and they provide a signal of satiety that has negative valence when strongly activated (Campos et al. 2016). The PBN has been characterised as a hedonic hotspot, the modulation of which by either GABA or Benzodiazepines potently modulates experienced reward (Söderpalm & Berridge 2000); ARC_AgRP_ neurons GABA-ergically inhibit PBN neurons, thus stimuli predicting glucose consumption should inhibit ARC_AgRP_, releasing the PBN from inhibition (Qunli Wu et al. 2014). Further, hormones related to hunger and feeding (GLP-1 & leptin) modulate PBN activity and subsequent behaviour (e.g. Alhadeff, Baird, et al. 2014; Alhadeff, Hayes, et al. 2015). Of note, these three valuation systems all project to and modulate the dopaminergic neurons in the ventral tegmental area(VTA_DA_). The interface between these hypothalamic-brainstem networks and the VTA_DA_, is arguably the most important interface for mediating the dialogue between energy homeostasis and value computation.

While most evidence for encoding of RPEs is obtained under homeostatic deprivation, the modulation of RPE signalling triggered by physiological fluctuations in glucose availability (glycemic flux) remains yet to be characterised in the human brain. This begs the questions, how are RPE signals modulated by these subcortical circuits that integrate, evaluate, and predict energy-homeostatic states? We hypothesize that glucose fluctuations above and below average levels of serum glucose, will down and up modulate RPE responses in regions of interest. To test these hypotheses, we acquired a large volume of fMRI data in five participants during a simple Pavlovian cue-conditioning task, while their serum glucose was systematically manipulated.

## Methods

### Subjects

Five healthy, normal-weight subjects in the age range 23 to 29, participated in the study. Exclusion criteria were: 20 > BMI > 25; 18 > Age > 32 yrs; any metabolic or endocrine diseases or gastrointestinal disorder; any known medication that might interfere with the study; claustrophobia; and any metal implants or devices that could not be removed. Informed consent was obtained from all subjects as approved by the Regional Ethics Committee of Region Hovedstaden (protocol H-4-2013-100) and in accordance with the declaration of Helsinki.

### Experimental procedure

The experimental design constituted a single-blinded, randomised control trial, with repeated measures crossover-design. On four separate days, subjects fasted for a minimum of ten hours before testing. At the beginning of an experimental session, subjects ingested either a hi-glucose (75 g, 300 kcal) or lo-glucose preload (10 g, 40 kcal) diluted to 100 ml with a 0-kcal lemon juice masking the taste of the glucose. Both preloads were anecdotally reported by independent samplers to be highly palatable.

### Experimental task

After consuming the preload, participants engaged in a simple pavlovian cue-conditioning task. The colour of the fixation cross cued both the onset of each trial (Cue_onset_), as well as stochastically predicting glucose delivery (Fig. 1a), with one colour signalling a high probability of glucose delivery (Cue_high_), and another signalling a low probability (Cue_low_). 10–15 seconds after delivery of oral stimulus, a purple cross signalled that subjects were allowed to swallow. All probabilities and contingencies were implicitly revealed only through experience in the scanner, and all were stationary over all test days. The mapping between colour and outcome probabilities was counterbalanced across subjects, while mapping was stationary within and between sessions. Participants went through ~82 trials [82 ± 1.5 SEM] each day giving ~328 trials per subject. Serum glucose measurements were attained immediately before and 20 minutes after ingestion, using a Contour^®^ Next glucose meter (Fig. 1b). As expected, prior to ingestion (t0) there was no significant difference between hi- or lo-glucose days [4.6 ± 0.4 SEM], whereas twenty minutes after ingestion (t20) there was [lo-glucose mean = 4.8, hi-glucose mean = 6.9].

**Figure 1.**
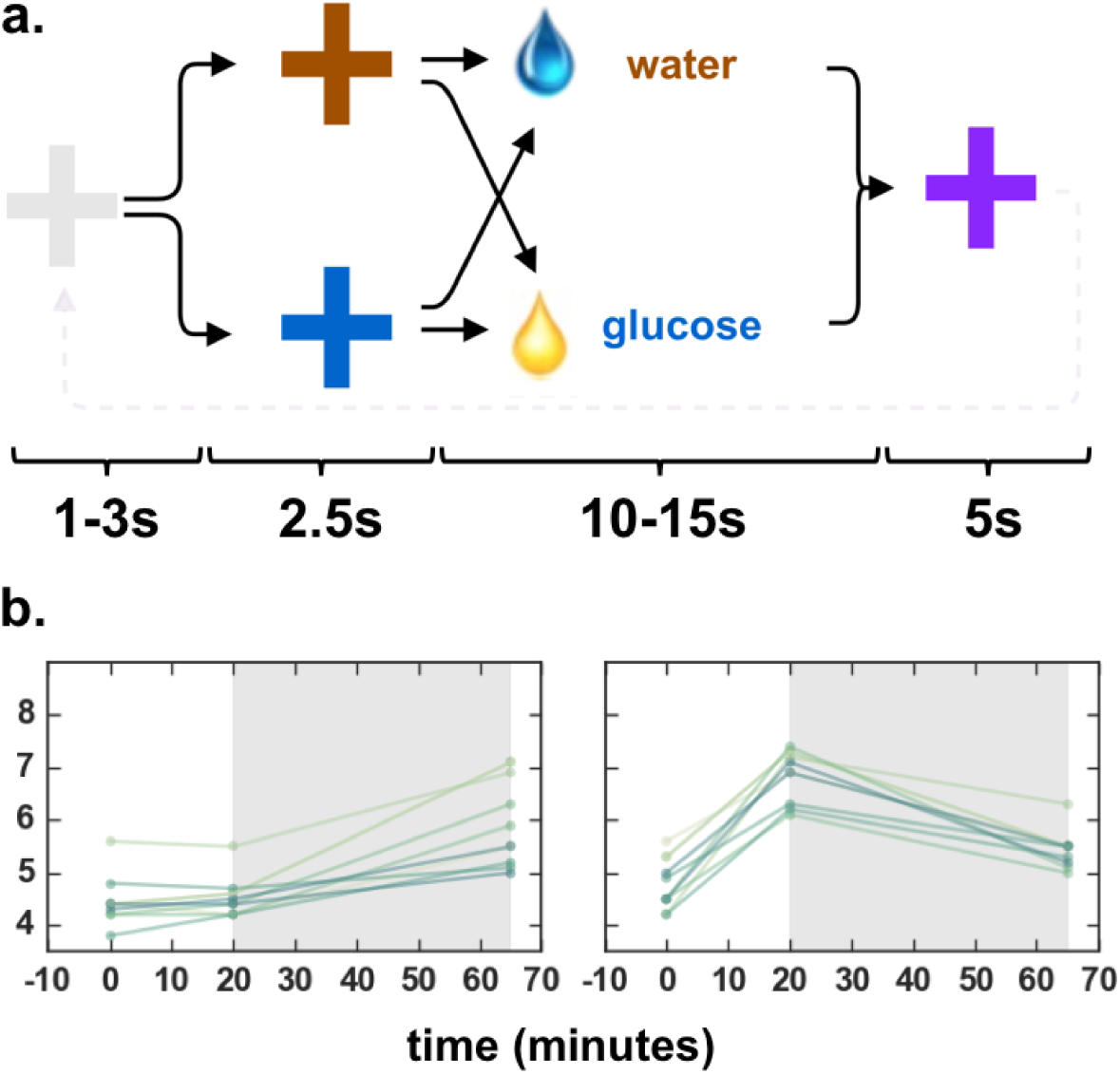
Experimental design and glucose trajectories. **a**, At Cue_onset_ participants are presented with a neutrally coloured fixation cross (grey) for 1–3s after which either Cue_high_ (here illustrated as blue cross) or Cue_low_ (brown cross) is presented with 0.5 probability each. Each cue signalled either high probability (0.8) of glucose delivery (0.4 ml) and low probability (0.2) of water delivery (0.4 ml), or the converse probabilities, respectively. A fixed duration after presentation of either cue (2.5 seconds), the liquid stimuli was delivered over 2.5 seconds. This was followed by 10–15s jitter and a cue for swallowing (purple) that lasted for 5s. after which a new trial initiated with the onset of the neutral cross. **b**, Plot of measured serum glucose (y-axis) over each session (x-axis) that lasted approximately 65 minutes. Left shows the lo-glucose preload sessions, while right shows hi-glucose preload sessions. The grey shading indicates the duration of the fMRI acquisition of a single session.

### Scanning procedure

Task related changes in regional brain activity were mapped with blood oxygen dependent (BOLD) MRI immediately after the second glucose measurement (t20). Functional MRI measurements were performed with a 3T Philips Achieva and a 32 channel receive head coil using a gradient echo T2* weighted echo-planar image (EPI) sequence with a repetition time of 2526 ms, and a flip-angle of 80°. Each volume consisted of 40 axial slices of 3 mm thickness and 3 mm in-plane resolution (220 × 220 mm). The axial field-of-view was 120 mm covering the whole brain, cutting off the medulla oblongata partially. During each session, 800 EPI volumes were acquired, resulting in 3200 EPI volumes per subject. Further, an anatomical T1-weighted image was recorded for each subject. Respiration and heart rate were measured to assess and model possible artefacts. Liquid tastants were contained in two 50 ml syringes, one containing water-only (water hence) the other containing glucose and water (glucose hence) solutions, attached to two programmable syringe pumps (AL1000-220, World Precision Instruments Ltd, Stevenage, UK), controlled by the stimulus paradigm script. The liquid was delivered orally via two separate 5m long 3mm wide silicone tubes. Each tube was attached to a gustatory manifold specifically built for the Philips head-coil (John B. Pierce Laboratory, Yale University). Visual stimuli were presented on a screen positioned 30 cm away from the scanner.

### Pre-processing

Pre-processing and image analysis were done using SPM12 software (Statistical Parametric Mapping, Wellcome Department of Imaging Neuroscience, Institute of Neurology, London, UK). To correct for motion, EPI scans were realigned to their mean using a two-step procedure and co-registered to the T1 weighted anatomical image through a unified segmentation procedure (Ashburner & Friston 2005). The realigned images were spatially normalised to the standard ICBM space template of European brains (Mazziotta et al. 1995), with a resampled voxel size of 3 mm.

### Modelling RPEs

At the first level, a general linear model (GLM) was set up to model cue and outcome related brain activity. We specified separate regressors which modelled the onset of Cue_onset_, Cue_high_ and Cue_low_ as well as outcome onsets for Outcome_gluc_ & Outcome_water_. Fig. 2a illustrates how the expectation value of glucose volume delivered evolves over time as a function of the cues observed. We specified RPE contrasts which were formulated by linear combinations of regressors, weighted as a function of the RPE values from the temporal-difference learning algorithm (Sutton & Barto 1998). As subjects learn the contingencies between visual stimuli (colour crosses) and outcome (juice or water) the RPE converge to the expected (average) value of the glucose content. This is conditioned on the cues that have been experienced and is illustrated in Fig. 2b. In this paradigm, there was no behaviour to fit a learning rate parameter to, so the steady-state values of the RPE was used instead. In effect, this assumes that the subjects learned the contingencies from the beginning. The effect of serum glucose on RPE was modelled by multiplying the resulting RPE by subject specific demeaned serum glucose (state hence), linearly interpolated between out-of-scan measurements. We specified the following contrasts of interest: RPE_pos_, RPE_neg_ with their state-weighted counterparts RPE_pos*state_, RPE_neg*state_ computed as first order parametric modulators.

**Figure 2.**
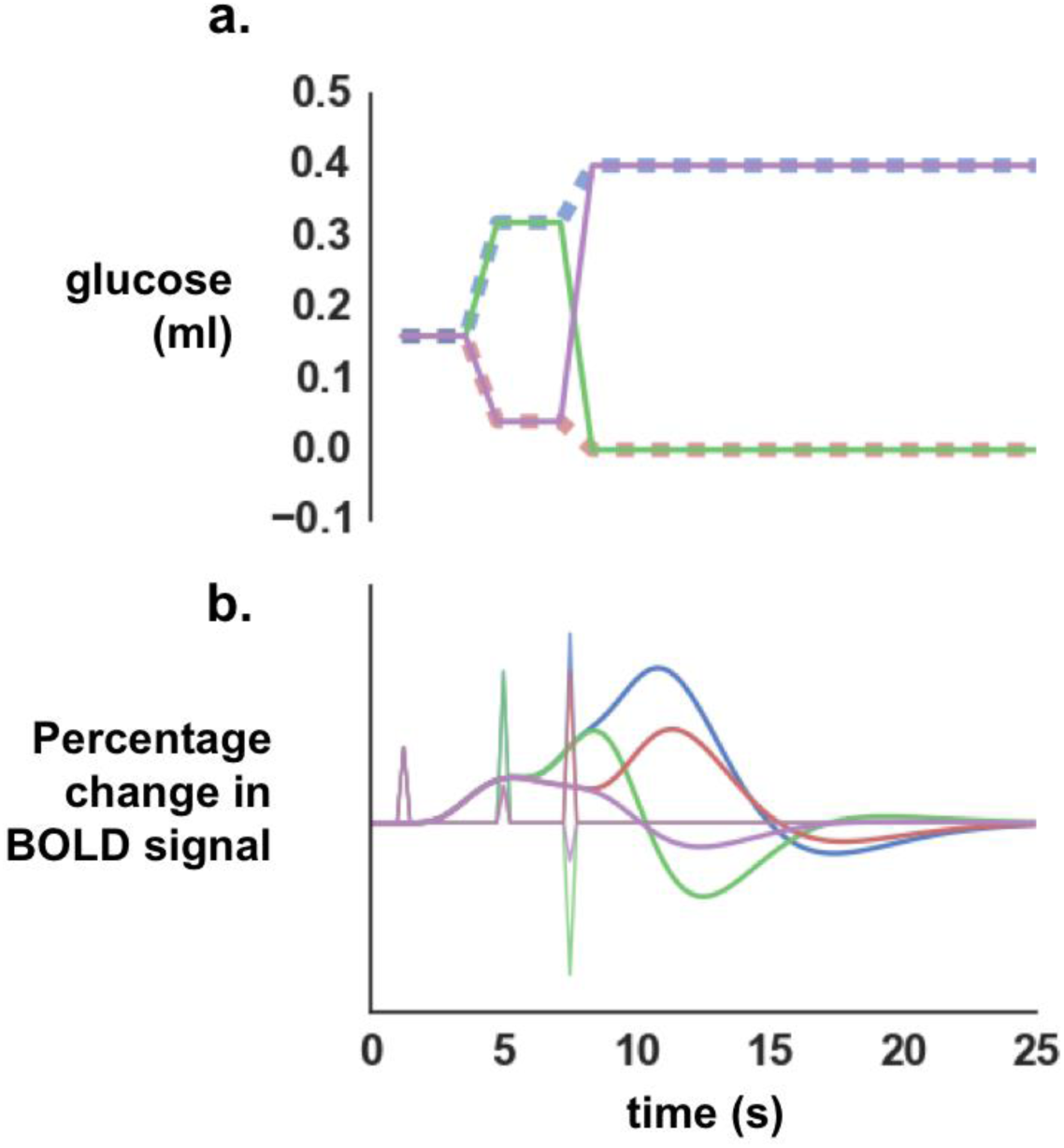
Expectations and fitted responses for reward prediction. **a**, Line graph depicts the objective reward expectations, expressed as the expected value in ml glucose, and the perturbation of these expectations under the onset of the experimental cues and outcomes. The dashed transparent lines illustrate when cues signalled high (or low) outcomes truthfully (blue high, orange low). The solid line illustrates when cues where paired with low-probability outcomes (green for high to low & purple from low to high). Note that reward expectations are updated three times: 1) at the onset of the Cue_trial_, 2) at the onset of Cue_high_ or Cue_low_, and 3) at the onset of Outcome_gluc_. or Outcome_water_. **b**, Illustrates simulated BOLD responses to RPE signals resulting from the updated reward expectations shown in Fig. 2a, generated by convolving the canonical hemodynamic response function with the RPE stick functions evoked by changes to the reward expectations with **<u>β = 1</u>**.

### fMRI analysis

After model specification, the sessions were concatenated using the function spm_fmri_concatenation (SPM 12) for each subject and a standard first-level fixed effects models was run over all subjects. All variables of interest were convolved with the canonical hemodynamic response function and fitted to the data using the specified GLM. The temporal evolution of cues and outcomes were modelled as separate conditions, each with state as parametric modulators. Regressors of no interest included a discrete cosine transform based 1/128 Hz cut-off frequency high-pass filter, rigid body realignment parameters using a 24 parameter Volterra expansion (Friston et al. 1996) and physiological noise from heart rate and respiration using the RETROICOR method {Glover:2000wy}. We specified the striatum (caudate, putamen and nucleus accumbens), brainstem (pons, ventral tegmental area and substantia nigra) and hypothalamus as Regions of interest (ROI). These ROIs were determined on basis of the literature describing dopamine projections from midbrain to the striatum and its role in regulating behaviour as a function of reward. The pons was selected to accommodate the literature described above, which sets certain nuclei within the pons as important homeostatic modulators. All ROI were defined with the WFU pick atlas (Lancaster et al. 2000; Lancaster et al. 1997). All initial first-level analysis was performed as whole-brain uncorrected at p < 0.001. Significant clusters in regions of interest (ROI) are all reported as small-volume corrected with a family-wise threshold of p < 0.05 at cluster level (abbreviated SVC FWE), unless otherwise stated.

## Results

### Cue induced brain activity

The “trial onset” cue signalled the expected value of glucose reward for the whole trial (Stauffer et al. 2014) and triggered an increase in activity in VTA bilaterally (Fig. 3a). Cue-induced VTA activation is consistent with existing evidence of VTA signalling RPE (e.g. D’Ardenne et al. 2008; Page et al. 2011; Eshel et al. 2016). The onset cue also led to deactivation of postcentral gyrus (primary somatosensory cortex), mediodorsal thalamus, and likewise in the striatum [whole brain, uncorrected p < 0.001] (not shown). In several brain regions, regional task-related activity changed in proportion with the magnitude of positive-going (i.e. better-than-expected) RPEs or negative-going (i.e. worse-than-expected) RPEs. Task related activity scaling with the RPE_pos_, formalized as an RPE-weighted linear combination of Cue_trial_, Cue_high_, and Outcome_gluc_, was found in left lateral caudate nucleus (Fig. 3b). Conversely, task related activity reflecting RPE_neg_, formalized an RPE-weighted linear combination of Cue_low_ and Outcome_water_, was located in the caudate nucleus bilaterally Fig. 3c), the medial dorsal thalamic nucleus, and insula.

**Figure 3.**
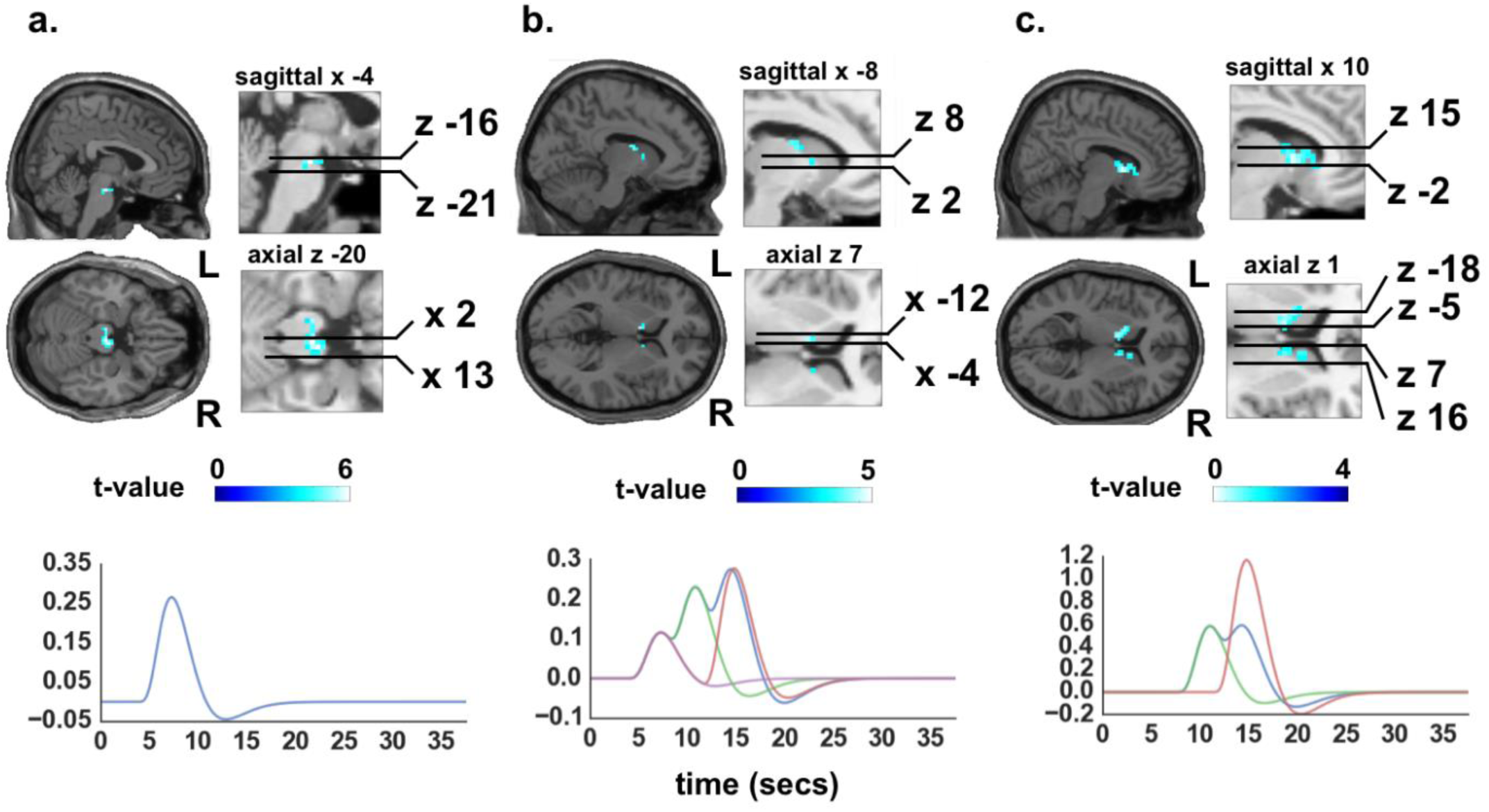
Statistical parametric maps of main effects of trial onset and RPE and fitted response. **a**, Main effect of trial onset cue, which reflects an RPE following the mean reward expectation for the whole trial, revealed activity in VTA bilaterally (*β* = 2.77) (R: [4 −17 −20] and L: [−8 −17 −20], FWE SVC). Further this revealed deactivation of precentral gyrus (primary somatosensory cortex), mediodorsal thalamus, and striatum (FWE whole brain, not shown). **b**, The main effect of RPE_pos_ revealed activity in left lateral caudate [*β* = 1.21; coordinates −8 4 7; FWE SVC]. **c**, The main effect of RPE_neg_ revealed bilateral activity in caudate (L: −11 −2 13; R: 10 7 1; *β* = 12.2] medial dorsal thalamic nucleus [7, −2, 22], and lateral insula [43, −2, −17] (all FWE). All fitted responses were generated by convolving the canonical hemodynamic response function with the RPE stick function multiplied by their respective beta-values extracted from the local maxima of the ROI.

### Modulation of task-related brain activity by homeostatic glycemic state

We were interested to identify changes in RPE processing over time as serum glucose either ascended or descended. A bilateral cluster, including the parabrachial nuclei (PBN), showed a modulation of the regional neural responses to positive RPEs by the glycemic state dynamics (Fig 4a). Higher levels of serum glucose amplified the response to RPE_pos_ in the PBN region (Fig. 4b). The main effect of RPE_neg*state_, which models the interaction between RPE_neg_ and state, did not yield any significant results in any ROI, or in exploratory analyses using uncorrected thresholds, in positive or negative contrasts. When considering both ascending and descending serum glucose fluctuations together, there was no detectable region where the RPE_neg_ signal was either positively or negatively modulated by serum glucose. Brain responses to the “onset cue” were also not altered by glycemic state dynamics.

**Figure 4.**
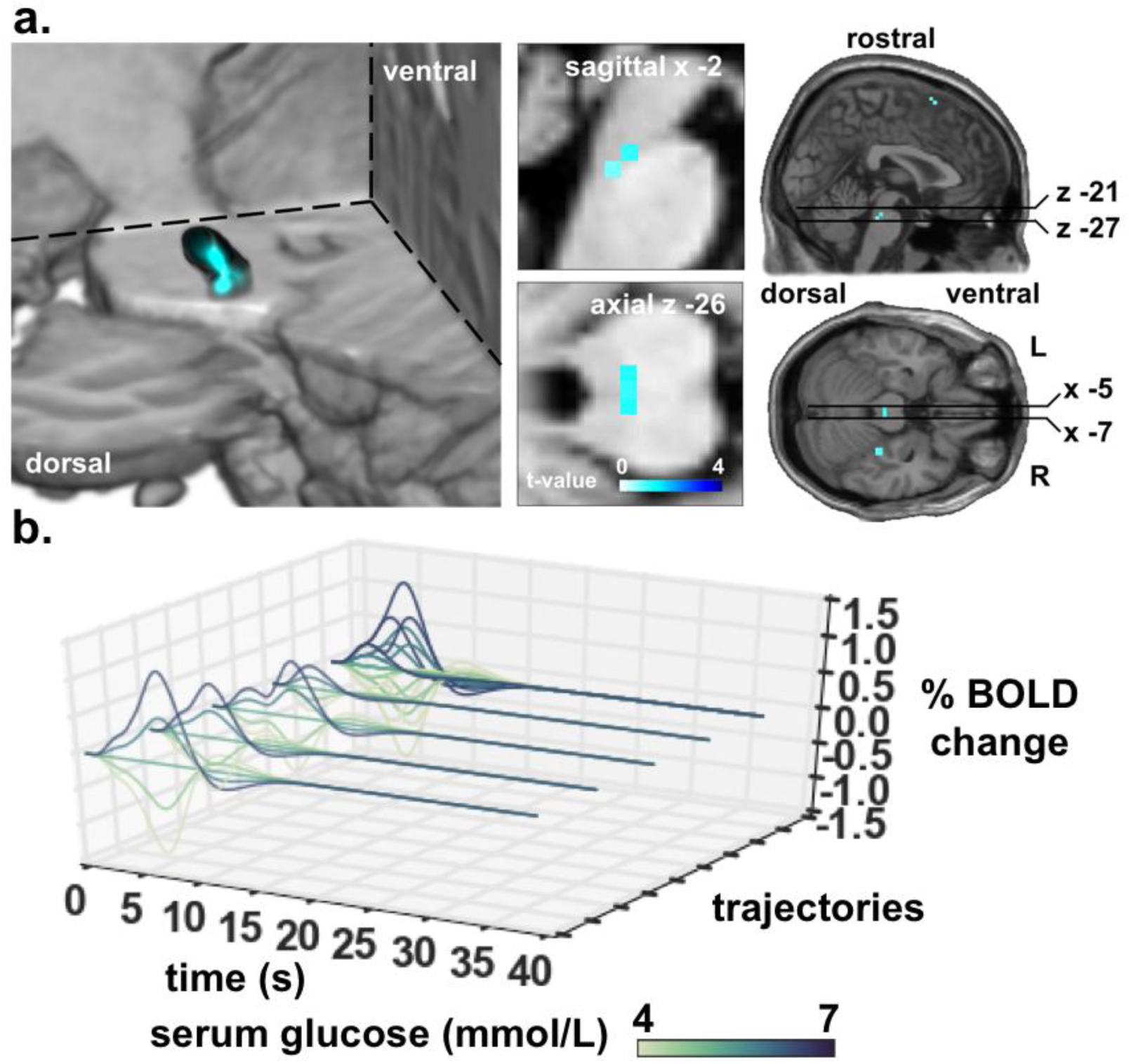
Statistical parametric maps of RPE_pos*state_ and fitted responses over varying glucose state. **a**, The main effect of RPE_pos*state_ revealed bilateral activity in the PBN [−2 −29 −26; FWE SVC]. **b**, Fitted response (*β* = 1.66) of the local maxima of PBN cluster (7 voxels) to the four possible trajectories that RPE_pos*state_, yield (see Fig. 2b) modulated by serum glucose state. The furthest trajectory on the y-axis are all four trajectories superimposed on to each other and is the signal which statistics is shown above in Fig. 2a.

We also tested for state-dependent modulatory effects on RPE processing which depends on whether serum glucose was ascending (Fig. 1b, left) or descending (Fig. 1b, right) over time. This yields four different contrasts (ascending vs. descending over RPE_pos*state_ and RPE_pos*state_) that are directly relevant to glucose state. Subtracting descending trajectories from ascending and vice versa, revealed no significant activity changes for RPE_pos*state_ [whole brain, uncorrected]. The same comparisons for RPE_neg*state_ did reveal significant effects in VTA and substantia nigra for ascending trajectories relative to descending trajectories (Fig. 5a). This result shows a relative amplification of the RPE_neg*state_ signal as glucose state increases. In instances where reward was lower-than-expected (thus yielding negative RPE), the glucose state modulated the RPE_neg_ signal in VTA and SN more so when glucose levels were ascending than descending.

**Figure 5.**
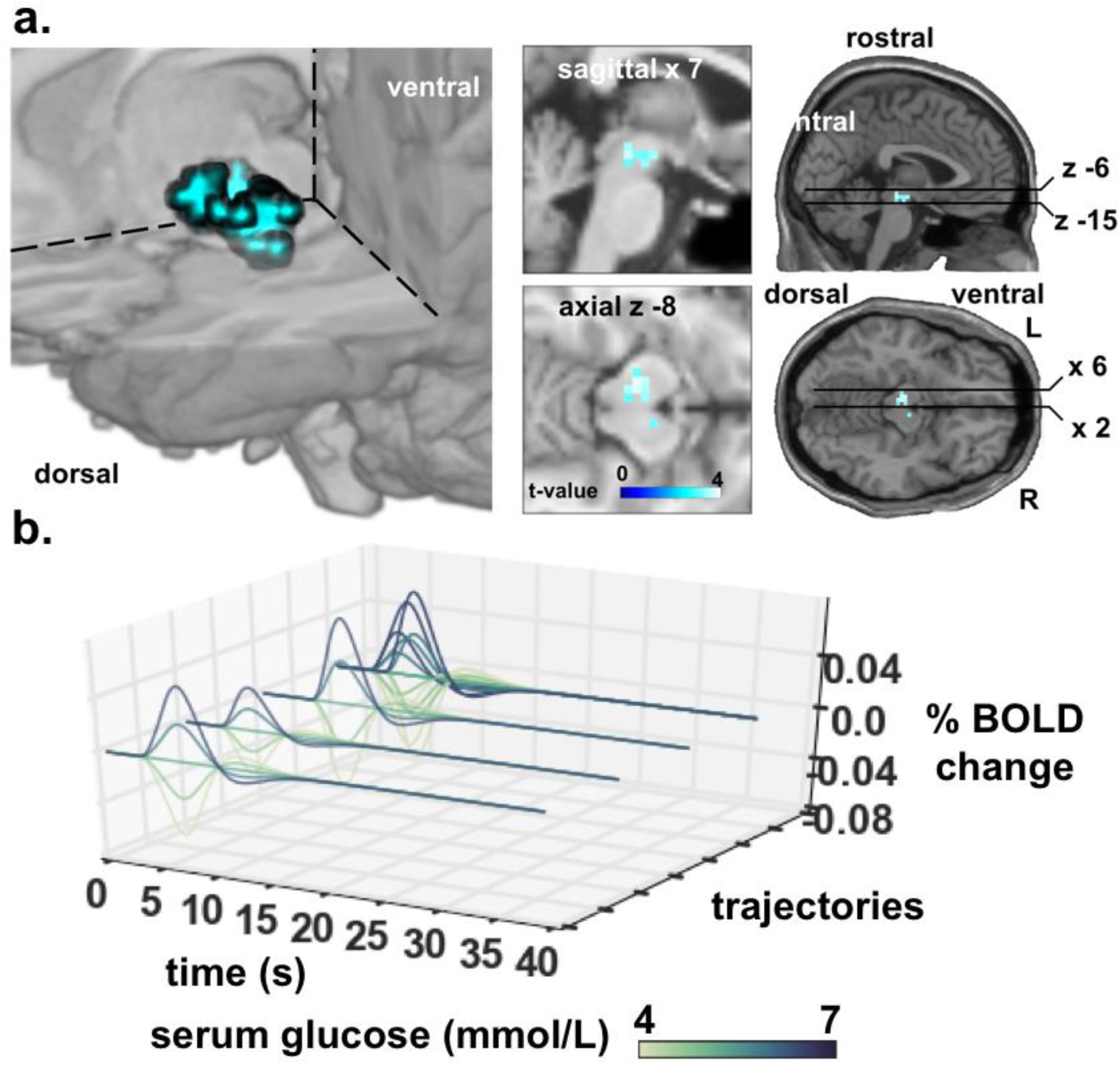
Statistical parametric maps of RPE_neg*state_ subtracted for increasing minus decreasing. **a)** Negative reward prediction error RPE_neg*state_ revealed glucose modulated activity in SN [±12, −22, −10] and VTA [0, −15, −6] when subtracting the effect of descending from the ascending glucose state [FWE SVC]. **b)** Fitted response (*β* = 0.34) of the local maxima of cluster [7, −11, 8; 52 voxels] to the three possible trajectories that RPE_neg*state_ yield modulated by serum glucose state. Onsets are not at zero because the negative trajectories do not envelop the trial mean which has a positive expectation.

## Discussion

We studied five individuals repeatedly with fMRI under increasing or decreasing levels of glucose, while participants performed a simple cue-conditioning task involving the probabilistic delivery of glucose or water in a single trial. Reward prediction error signalling in the parabrachial nuclei scaled positively with serum glucose levels during ascending and descending glycemic trajectories. The VTA and SN became more sensitive to negative RPEs for ascending compared to descending glycemic trajectories. We begin by discussing the interpretation of these state-modulated RPE effects, before considering other effects, and the limitations inherent under this paradigm.

In rodent models, the PBN acts as a 2^nd^ order relay of inputs from the nucleus tractus solitarius, and is critical in the control of energy homeostasis via its projections to amygdala (Norgren 1978; Loewy 1998), VTA (Miller et al. 2011), hypothalamus (Norgren 1976; Loewy 1998) and the nucleus accumbens (Li et al. 2012). Subnuclei of the PBN are targeted by descending projections from several nuclei implicated in energy homeostasis, including hypothalamus, amygdala, and the bed nucleus of the stria terminalis (Zhang et al. 2011; Loewy 1998). The PBN is known to be a potent site of reward modulation and subsequent behaviour in rodents. Microinjection of benzodiazepines (Söderpalm & Berridge 2000; Qi Wu et al. 2009; De Oliveira et al. 2011), endocannabinoids (DiPatrizio & Simansky 2008), opioids (Wilson et al. 2003; Chaijale et al. 2013) and melanocortin agonists (Skibicka & Grill 2009) into the PBN, all evoke hyperphagia s. To the best of our knowledge, the involvement of PBN in context of hedonics and reward signalling in the human brain remains yet to be charted. Here we provide novel evidence that PBN activity generates a gluco-sensory scaled positive RPE signal which is time-locked to both the sensory cues predicting glucose, as well as glucose outcomes.

Unlike the state modulation of serum glucose trajectories on the RPE_pos_ signal, we found no general state modulation of RPE_neg_ signalling, expressed during ascending and descending glycemic trajectories. Here the modulatory effect of the glycemic trajectory depended on whether glucose trajectories were ascending or descending. Regional activity scaling with RPE_neg_, the VTA and SN showed significantly higher state modulation effects during ascending vs. descending glycemic paths. In our experiment, the ascending glucose trajectory resulted from a low-glucose preload with the subsequent increase over time likely occurring by virtue of the continual ingestion of glucose throughout the paradigm (Fig. 1b). In the ascending condition, the neural response to RPE_neg_ is attenuated at lower levels of serum glucose, while it becomes amplified by the transition to higher serum glucose. Given that dopaminergic neurons of the VTA and SN are directly inhibited by insulin (Palmiter 2007), it is likely that the insulin release following hi-glucose preload was highest at the start of the paradigm, decreasing over time, and thus resulting in a gradual decrease in inhibition. The difference in RPE_neg_ in its state modulation between ascending and descending may therefore be attributed to differential dynamics of insulin secretion (see (Sun et al. 2014), though other hormones such as ghrelin (Malik et al. 2008; Kroemer et al. 2013; Sun et al. 2014) or leptin (Domingos et al. 2011; Figlewicz et al. 2003; Fulton 2000; Alhadeff, Hayes, et al. 2014; Takahashi & R. D. Cone 2005) may play a role.

Our finding that the VTA and SN responses are linked to RPE_neg_ may appear counterintuitive, given that these midbrain regions are typically associated with BOLD responses signalling positive-going RPEs. This is assumed to be by virtue of the fact that a greater range of firing rates can be devoted to the better-than-expected range, signalled by above baseline firing. This is contrasted to the worse-than-expected range, which can only be signalled by a decrease from an already low baseline frequency. It is conceivable that what we are asserting as being RPE_neg_ is in fact a positive RPE resulting from the gradual avoidance of glucose, which increases in magnitude with increasing levels of serum glucose as reported in humans (Cabanac 1971) and rats (Berridge 1991). Thus, as the experimental paradigm continues, especially under the conditions of glucose preload, serum glucose increases, and this may change the valence of the outcome, switching the affective connotation of glucose from palatable to aversive.

As detailed in the introduction, little is known about the principles how the interface between dopaminergic RPE signalling and energy homeostasis is implemented in the human brain. While there are many means by which circulating factors can modulate activity in the VTA and SN, the mechanisms by which this is mediated cannot be revealed without wider hormonal assays. Contemporaneous hormonal sampling, as well as continuous glucose monitoring in the scanner will prove an important step in revealing these hidden mediating factors.

From a theoretical perspective, results as presented here could be predicted by any model that invokes the notion of RPEs in service of homeostatic regulation. For example, models inspired by optimal control theory such as Homeostatic Reinforcement Learning ((Keramati & Gutkin 2014) or MOTIVATOR theory (Dranias et al. 2008). Alternatively, under the theory of Active Inference, phasic dopamine is recast as encoding updates to the precision assigned to the behavioral policies that lead to desired outcomes, that (in this context) remediate long-run homeostatic error (Schwartenbeck et al. 2015).

There are several technical limitations that should be noted in discussing this experiment. Though relatively high volumes of functional data (150 minutes per subject) were acquired in each subject, the total number of subjects was low. Future work will expand this paradigm with a larger group of to afford random effects modelling, and thus generalisation to the population sampled from. In contrast to our hypotheses, we found no modulatory effect of hypothalamic nuclei on RPE signalling. We would like to stress that the current imaging protocols and field-strength (3T) were not optimal to dissociate neural activity in the hypothalamic nuclei. Due to the proximity of air sinuses adjacent to the hypothalamus and the effective resolution available, the present study most likely had insufficient sensitivity to capture activity in hypothalamic regions of interest. Future work at higher field strengths (7T) may overcome these limitations. Finally, the cue-conditioning employed in this study was passive. Hence, subjects produced no overt choice behaviour against which to fit learning rate parameters for the RPE model, instead we relied on the asymptote values for the RPE signals. The problem of modelling RPEs in the absence of choice behaviour, motivates fitting learning rate parameters directly to brain data, a computational imaging approach that future work will exploit (Meder et al. 2017)

In conclusion, we exploited a simple paradigm, capable of eliciting RPEs under differential glycemic trajectories, to identify brain stem structures that show a modulation of RPE signalling dependning on the glycemic homeostatic state. We found that the PBN signals a positive-going reward prediction that is subject to systematic modulation by serum glucose. In the VTA and SN, negative-going RPEs were modulated by serum glucose trajectories, but in a way that was specific to an ascending glycemic slope. Together the results show that RPE signals in key brainstem structures are modulated by homeostatic trajectories inherent in naturally occurring glycemic flux, revealing a tight interplay between homeostatic state and the neural processing of primary reward in the human brain.

## Acknowledgements

We thank Mehdi Keramati and Boris Gutkin for several helpful discussions. This work was supported by the following funders: H.R.S (Lundbeck Foundation Grant of Excellence “ContAct” ref: R59 A5399; Novo Nordisk Foundation Interdisciplinary Synergy Programme Grant “BASICS” ref: NNF14OC0011413) O.J.H (Lundbeck Foundation, ref: R140-2013-13057; Danish Research Council ref: 12-126925) T.M (Lundbeck Foundation ref: R140-2013-13057).

## Author contributions

T.M. & O.H conceived of the study, and all authors designed the study. T.M collected the data and visualised the results. All authors contributed to analysis and interpretation of data, and to the writing of the manuscript.

